# FlavoTyper: a genome-based *in-silico* serotyping tool for the fish pathogen *Flavobacterium psychrophilum*

**DOI:** 10.64898/2026.09.14.751350

**Authors:** Salma Mbarki, Pavla Debeljak, Mathilde Carpentier, Keith A. Jolley, Noëlle Haddad, Tatiana Rochat, Eric Duchaud

**Author notes:** **Corresponding author and email address** Eric Duchaud. **Repositories** Newly sequenced *F. psychrophilum* genomes have been deposited at the ENA database BioProject PRJEB21067 as well as in the PubMLST database.

## Abstract

*Flavobacterium psychrophilum* is a devastating pathogen of fish reared in freshwater worldwide. Serotyping is a relevant method for epidemiological surveillance and outbreak detection, as well as for a better understanding of host-pathogen interactions. Serological diversity may also have important consequences for the selection of appropriate strains for vaccine development and for selective breeding for increased disease resistance. *F. psychrophilum* serotyping relies on structural variations in the O-polysaccharide (O-PS) moiety of the cell surface lipopolysaccharide (LPS). However, conventional serotyping is costly, labor-intensive and requires significant technical expertise. Moreover, divergent scheme proposals highlighted the absence of harmonization among laboratories. In this context, the development of an mPCR-based serotyping scheme targeting *wzy* genes greatly improved the reliability and standardization of serotyping. Nevertheless, the proposed mPCR scheme did not capture the entire diversity of genomic variability. The aim of this study was to establish a robust and publicly available tool for *F. psychrophilum* genome-based serotyping. Extensive genome analysis of the O-antigen biosynthesis locus allowed the identification of biomarkers enabling the development of FlavoTyper, an *in-silico*-based serotyping tool. The FlavoTyper tool was evaluated on all *F. psychrophilum* genome assemblies publicly available, providing sound and sensitive predictions and easily interpretable results. When applied to a curated collection of publicly available genomes, the *in-silico* O-types assigned by the tool were statistically significantly associated with host fish species, confirming previous studies — coho salmon with O:0, rainbow trout with O:1 and O:2, and ayu with O:3 — and their distribution across MLST clonal complexes revealed that the O-antigen locus is frequently rearranged independently of the core-genome lineage, consistent with the extensive recombination that shapes the evolution and genomic diversity of this species.

**Impact statement:** We developed FlavoTyper, the first genome-based *in-silico* serotyping tool for *F. psychrophilum*. FlavoTyper accepts *F. psychrophilum* genome assemblies in FASTA format and both single and multi-genome runs are supported. The pipeline includes quality control checks of the input data and the reported results are readily interpreted by the users with clear warnings when required as well as a locus map. Applied to publicly available genomes, FlavoTyper allowed the identification of 23 distinct serotypes and captured the host associations established over three decades by conventional and PCR-based methods. Distributed openly through PyPI, Bioconda and Galaxy, also as a plugin in the Bacterial Isolate Genome Sequence Database (BIGSdb) of the PubMLST platform, the tool provides researchers and veterinary professionals in aquatic animal health with a reproducible, scalable, and standardized framework for epidemiological surveillance and vaccine development for *F. psychrophilum*.

**Data summary:** The authors confirm all supporting data, code and protocols have been provided within the article or through supplementary data files.

Newly sequenced genomes have been deposited under EBI BioProject PRJEB21067.

(1) The FlavoTyper source code and database files are accessible from the GitLab repository https://forge.inrae.fr/eric.duchaud/flavotyper, achieved on Zenodo: https://doi.org/10.5281/zenodo.22259350, and installable through conda and pypi.
(2) The tool is also available as a Web-based interface through Galaxy Europe ([FlavoTyper/Galaxy Europe](https://usegalaxy.eu/?tool_id=toolshed.g2.bx.psu.edu%2Frepos%2Fiuc%2Fflavotyper%2Fflavotyper%2F0.5.1%2Bgalaxy0&version=latest)). FlavoTyper has also been included in PubMLST allowing both *in-silico* MLST and serotyping from genome assemblies (https://bigsdb.readthedocs.io/en/latest/data_analysis/flavotyper.html).

## Introduction

*Flavobacterium psychrophilum*, from the *Flavobacteriaceae* family, within the phylum Bacteroidota (1), is currently one of the most devastating bacterial freshwater fish pathogens. It is the causative agent of rainbow trout fry syndrome and bacterial cold-water disease in salmonid fish worldwide (2). All salmonid species are believed to be susceptible to infection (3–5), but this bacterium also significantly affects wild and farmed ayu (*Plecoglossus altivelis*) (6). More recently, *F. psychrophilum* has been retrieved from moribund wild European eel (*Anguilla anguilla*) (7) as well as from different carp species including crucian carp (*Carassius auratus gibelio*), grass carp (*Ctenopharyngodon idella*) and silver carp (*Hypophthalmichthys molitrix*) suffering severe mass mortalities (8,9). *Flavobacterium psychrophilum* is therefore of serious concern for aquaculture and wild fish populations.

Pathogen typing is essential for effective surveillance and outbreak detection (10 for recent review). In the last decades, different molecular typing schemes have been proposed for *F. psychrophilum* (11–14). Among those, MLST (15) – a core-genome based typing method – has been extensively used providing invaluable knowledge on the population structure and evolution at different times and geographic scales (5 and references therein).

On the other hand, serotyping relies on the antigenic diversity of surface-exposed structures such as capsular polysaccharides, lipopolysaccharides and surface proteins, which are frequently encoded by accessory genomic regions. These structures contribute substantially to bacterial virulence through their involvement in immune evasion, serum resistance, host-cell adhesion, inflammation and biofilm formation (16 for review).

Serological diversity among *F. psychrophilum* isolates was documented from the earliest studies, resulting in the proposal of different serotyping schemes and classifications. In Japan, Wakabayashi and colleagues (17) described two serotypes (O-1 and O-2), to which O-3 and O-4 were subsequently added (18,19). Meanwhile, Lorenzen and Olesen (20) in Denmark developed a distinct serotyping system based on three serogroups (Fp, Th, and FpT), whereas in Spain, Mata and colleagues (21) proposed a scheme able to identify as much as seven different serovars. These parallel approaches highlighted the absence of a unified serotyping framework. In addition, the conventional protocols currently available for serotyping *F. psychrophilum* isolates present several limitations, including the requirement for animal-derived antisera and the subjective interpretation of agglutination reactions. Moreover, some strains exhibit auto-agglutination, fail to react with available antisera, or cross-react with multiple antisera, necessitating labor-intensive reciprocal absorption procedures using heterologous strains. Furthermore, discrepancies between agglutination and ELISA results have been reported (20). Consequently, conventional serological serotyping remains costly, time-consuming, and highly dependent on specialized expertise.

In 2017, Rochat and colleagues (22) identified key molecular determinants of the serotypes and proposed a mPCR serotyping scheme allowing the identification of 4 types: 1, 2, 3, and 0 (the latter grouping all remaining undefined types). An additional primer pair was included in 2020 (23) allowing the identification of the type 4 serotype. This unified scheme was since applied to about 850 isolates retrieved from various fish species from worldwide origin (5 and reference therein). In addition, significant associations between *F. psychrophilum* molecular serogroup and host fish species were identified (5,22).

In 2019, Cisar and colleagues (24) formally demonstrated that the molecular determinants previously identified (22) encoded for the O-antigen (O-Ag) polymerase Wzy. According to the *wzy* gene type, the glycosidic linkage between d-Qui2NAc4NR and L-Rha varies: β(1–3) for wzy1 and α(1–2) for *wzy2*. Using a mouse monoclonal antibody against *F. psychrophilum* strain CSF259-93 (harboring *wzy2*) targeting the O-Ag and molecular dynamic simulations they pointed out structural differences in O-Ag conformations according to the nature of the produced linkage bond (25). In addition, they attributed the biosynthesis and/or transfer of the R1 group (*i.e.*, 3,5-dihydroxyhexanoyl) in d-Qui2NAc4NR1 to the presence of 5 genes (*wfpC, D, E, F and G*) encompassed in the O-Ag encoding locus. They also observed differences among strains in this latter group of genes, which is present in some strains and substituted by *wfpH* or *wfpI* in others, suggesting replacement of the R1 group with an unknown acyl group. In addition, R-group variability contributes independently to O-Ag antigenicity (24,25) suggesting that each O-Ag genomic structure represents an antigenically distinct O-serotype.

While the mPCR approach overcomes many limitations of serum-based serotyping, it classifies isolates into only five types and does not resolve the R-group variability.

The accumulation of whole-genome sequence (WGS) data has driven a profound methodological shift in bacterial typing and diverse *in-silico* tools have been developed for predicting serotypes from WGS data. Most of the proposed tools rely on the identification of suitable biomarkers and blast interrogation of a query genome against an accurate and comprehensive biomarker database. Ideal biomarkers for predicting O-Ag types are generally genes with key roles in determining the O-Ag structure. Despite the rapid expansion of *in-silico* serotyping tools (*e.g.*, SerotypeFinder (26), ECTyper (27), Kaptive (28), no equivalent is yet available for *F. psychrophilum*.

In the present study, we used extensive genome comparisons to identify biomarkers and developed FlavoTyper, an *in-silico* tool for *F. psychrophilum* serotyping from WGS data.

## Methods

### Genomic data resources

Two complementary genomic resources were exploited during this work. The first resource is composed of the complete set of *F. psychrophilum* assemblies available in NCBI GenBank which was retrieved on March 10^th^ 2026, yielding 482 assemblies with associated metadata (assembly statistics, BioSample descriptors and the NCBI ANI taxonomy-check result). This NCBI genome collection was used to evaluate the FlavoTyper tool. A second collection of 357 *F. psychrophilum* genomes was retrieved on May the 29^th^ 2026 from the PubMLST database (29), each accompanied by curated metadata (isolate name, country and continent of origin, host fish species, year of isolation and MLST clonal complex). This database had been manually curated to avoid duplicated genomes and was used for analyzing associations between serotype, host-fish species and clonal complex affiliation.

### Genome Comparisons

Genome comparisons were performed as previously described (22) using the web interface MicroScope (30). The MicroScope platform allowed an expert-curated annotation of *F. psychrophilum* genome assemblies, together with integrated tools for synteny analysis, gene-content comparison and pairwise genome alignment. This platform was used for the comparative-genomic analysis underpinning the FlavoTyper biomarker database. Gene organization analysis was conducted using synteny (*i.e.*, orthologous gene set computed using BlastP Bidirectional Best Hit or at least 30% identity on 80% of the shortest sequence and having the same local organization in two strains - gap of 5 genes tolerated). This work yielded the inventory of O-Ag locus architectures and allowed the identification of marker genes that underpin the typing scheme proposed in this work.

### Design of the *in-silico* serotyping tool

Boundaries of the biomarker genes or group of genes were manually delineated after pairwise sequence alignments. Special attention was given to the R1 variants, in which obvious intragenic recombination events were detected. Typing rules including sequence identity and % coverage thresholds that formally translate the biomarkers sequences into a serotype were computed after extensive pairwise comparisons. Alignments between the different Wzy protein types were performed using Pairwise Sequence Alignment from the EMBL-EBI Job Dispatcher sequence analysis tool (31). Three-dimensional models were generated with ΑlphaFold 3 (https://alphafoldserver.com/) (32), and model confidence was assessed using template model (pTM) scores. Structural similarity searches were conducted using DALI (33) and Foldseek (34).

### Implementation of the FlavoTyper software

FlavoTyper was developed and tested under Python 3.11 and requires Python 3.10 or any later version. Development was tracked with Git on the INRAE institutional GitLab forge. The tool relied on two external bioinformatics dependencies. BLAST+ (35) provides the alignment engine: the makeblastdb command builds searchable nucleotide databases from the reference biomarkers, and the BLASTn command performs all marker-versus-genome alignments. FastANI (36) was used as a first layer to verify that the input genome belongs to the species *F. psychrophilum*, by estimating the average nucleotide identity between the query and the *F. psychrophilum* type strain NCIMB 1947^T^; the default species-delineation threshold was set at ANI ≥ 95 %. FlavoTyper was released as a free and open-source software under the Apache-2.0 license; it is distributed through PyPI (pip install flavotyper), Bioconda (conda install -c bioconda flavotyper), the Galaxy Europe platform and also as a plugin in the Bacterial Isolate Genome Sequence Database (BIGSdb) of the PubMLST platform.

FlavoTyper is organised as a modular pipeline that accepts one or several genome assemblies in FASTA format and executes three modules sequentially (Figure 1): an input-and-quality-control module, a typing module, and an optional locus-analysis module that needs to be enabled by the user (--locus-analysis). The quality-control module performs a species-validation step during which each assembly is compared to the *F. psychrophilum* type strain NCIMB 1947^T^ with fastANI, and if the best ANI value of the input assembly falls below the threshold (default 95 %), the tool does not proceed with the typing. All details regarding the quality checks performed by FlavoTyper are provided in Supp. Table 1. The second, typing, module aligns the query genome against the reference biomarkers using BLASTn and assigns the serotype based on the detected markers. Different result categories may be returned following this step depending on the scenarios outlined in Supp. Table 2. The third, optional locus-analysis module aligns the reference O-Ag loci against the query genome, extracts the best-matching locus sequence in FASTA format, generates a pairwise sequence alignment and produces a map of the locus that highlights the detected biomarkers. The outputs produced by FlavoTyper for every run are summarized in Supp. Table 3.

**Figure 1:**
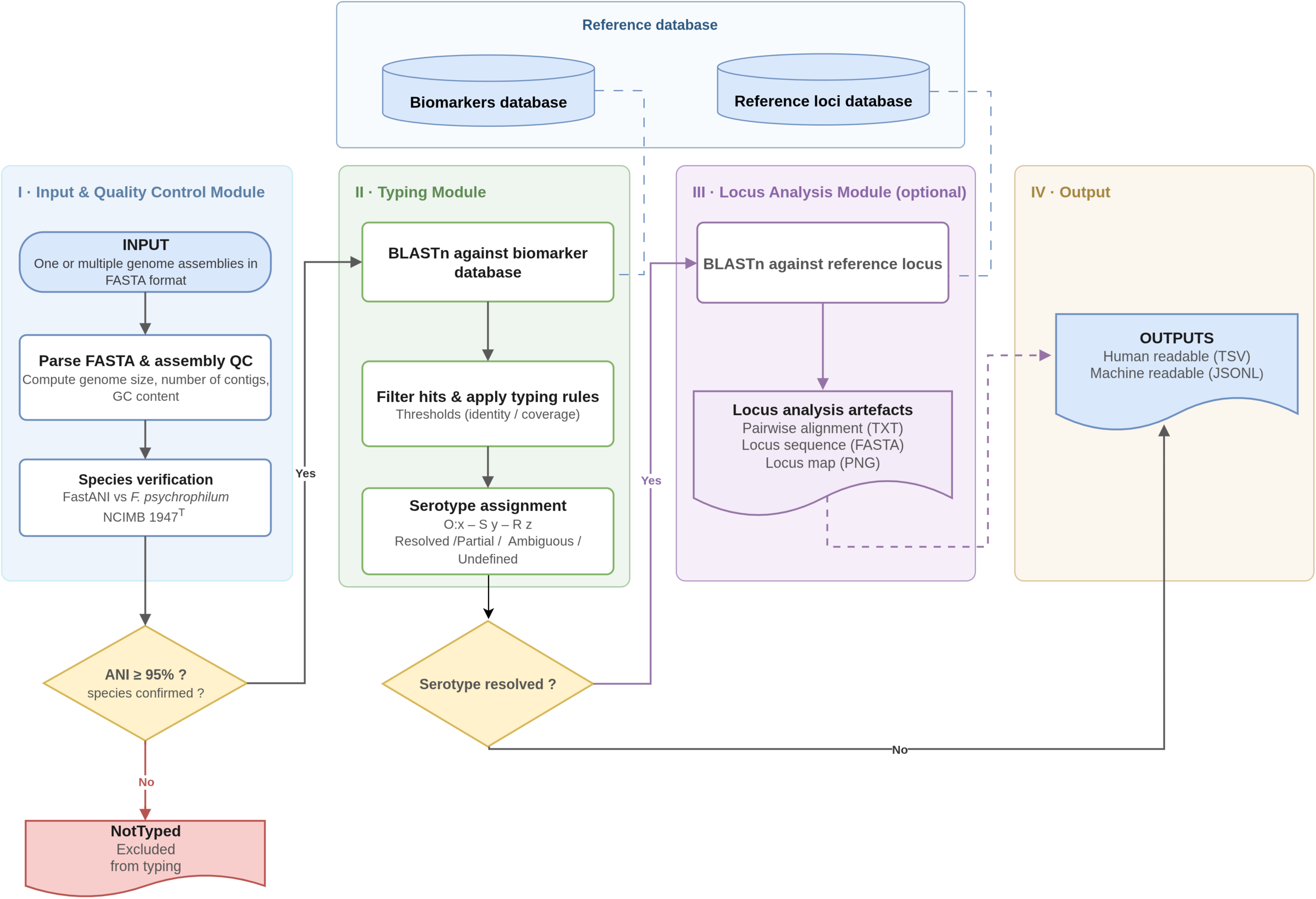
Flowchart outlining the major stages within FlavoTyper. Input can be either a single or multiple genomes. The first module corresponds to a species verification step coupled to a quality control of the input genome assembly(ies). The tool proceeds with the prediction of the serotype only when the species is confirmed to be *F. psychrophilum*. The second module is the typing *per se* through an alignment of the input genome(s) using BLASTn against the biomarker database, then the serotype is assigned based on the detected markers. A third and optional module allows the extraction of the O-Ag locus detected in the input genome(s) and plots a representative map with the genomic coordinates of the detected marker genes.

**Table 1:** Marker genes and their respective reference sequences integrated in the FlavoTyper biomarker database.

| Type | Marker gene(s) | Reference genome | Locus_tag | Length (nt) |
| --- | --- | --- | --- | --- |
| **O-type** |  |  |  |  |
| **O:0** | *wzy0* | OSU THC0290 | THC0290_2043 | 1,485 |
| **O:1** | *wzy1* | DK001 | DK001_50101 | 1,284 |
| **O:2** | *wzy2* | DK002 | DK002_320117 | 1,275 |
| **O:3** | *wzy3* | PH209 | PH0209_2216 | 1,317 |
| **O:4** | *wzy4* | DK150 | DK150_550174 | 1,326 |
| **O:5** | *wzy5* | FI146 | FI146_160075 | 1,008 |
| **O:6** | *wzy6* | KU220506-6 | KU2205066_40100 | 1,434 |
| **O:7** | *wzy7* | DK095 | DK095_460107 | 1,248 |
| **S-type** |  |  |  |  |
| **S1** | *wfpM,N,O,P* | LVDJ XP189 | LVDJXP189_340008 to<br>LVDJXP189_340005 | 3,299 |
| **R-type** |  |  |  |  |
| **R1V1** | *wfpC,D,E*<br>(*r1_core*) | DK001 | DK001_50109 to<br>DK001_50111 | 2,303 |
|  | *wfpF*<br>(*v1_specific*) | DK001 | DK001_50112 | 729 |
| **R1V2** | *wfpC,D,E*<br>(*r1_core*) | DK001 | DK001_50109 to<br>DK001_50111 | 2,303 |
|  | Rieske + *wfpF'*<br>(*v2_specific*) | PH209 | PH0209_2205 to<br>PH0209_2204 | 1,086 + 729 |
| **R1V3** | *wfpC,D,E*<br>(*r1_core*) | DK001 | DK001_50109 to<br>DK001_50111 | 2,303 |
|  | *wfpF''*<br>(*v3_specific*) | LM09a | LM09A_201010 | 744 |
| **R2** | *wfpH* | KU051128_10 | KU05112810_320053 | 642 |
| **R3** | *wfpI* | KU061128_01 | KU06112801_120028 | 618 |
| **R4** | *wfpJ,K,L* | JIP02/86 | FP1282 to FP1280 | 2,578 |

The *r1\_core* sequence (genes *wfpC*; *wfpD*; *wfpE*) is shared by all three R1 sub-variants. R0 and S0 correspond to the absence of R- or S-group markers and have no associated sequence.

### Core-genome phylogeny of *F. psychrophilum*

A whole-genome phylogeny was reconstructed for 334 *F. psychrophilum* genomes retrieved from the 357 of PubMLST for which the serotype has been resolved with FlavoTyper and using *F. columnare* strain F2S17 as an outgroup (335 genomes in total). All genomes were annotated with Prokka v1.15.6 (37) and the pan-genome was inferred with Panaroo (38) in strict cleaning mode (--core_threshold 0.98). Gene families that were single-copy and present in all 334 *F. psychrophilum* genomes were retained as the core set (959 families), and the corresponding *F. columnare* orthologue was assigned to each family by reciprocal best BLASTP hits (E-value < 1 × 10⁻⁵); the 869 families with a reciprocal outgroup orthologue were kept for phylogenetic analysis. Each family was aligned individually with MAFFT (39), trimmed with Gblocks (40), and the trimmed alignments were concatenated into a super matrix of 254,073 amino-acid positions. A maximum-likelihood tree was inferred with IQ-TREE 3 (41) under the LG+F+G4 substitution model. The tree was rooted on *F. columnare* and visualized with iTOL (42), onto which the O-, R- and S-type as well as the clonal complexes they belong to were mapped.

### Statistical Analysis

Associations between the *in-silico* O:x-type and host fish species (*Oncorhynchus mykiss*, *Plecoglossus altivelis*, *O. kisutch*, *O. tshawytscha* and *Salmo trutta*) were analyzed via Fisher’s exact test with Monte Carlo simulation (10⁵ replicates) using R version 4.2.2. The O:6 type was excluded from the analysis due to low sample size. Standardized Pearson residuals were calculated post-hoc and the Holm–Bonferroni method was applied to determine which species– O:x-type associations were significant (α = 0.05).

## Results and discussion

### Construction of the FlavoTyper biomarkers database

In 2017, from 34 *F. psychrophilum* genome assemblies, Rochat and colleagues (22) reported 15 different genetic structures for the O-Ag biosynthesis locus. In 2019, Cisar and colleagues mentioned 20 O-Ag genetic types among the 379 genomes available in GenBank at that time. We used the same path in order to identify additional genetic variations. Because *wzy1* and *wzy2* have been functionally characterized and used to develop the first mPCR typing scheme, we defined groups according to their *wzy* or *wzy*-like gene content (Supp. Figure 1). The *F. psychrophilum* O-Ag encoding loci begin with 8 genes and end with 5 genes conserved in all strains. Between these two groups of genes, lies a variable region encompassing 10 to 18 genes. Within this variable region, three loci were selected defining the O-type, the S-type, and the R-type (Figure 2).

**Figure 2:**
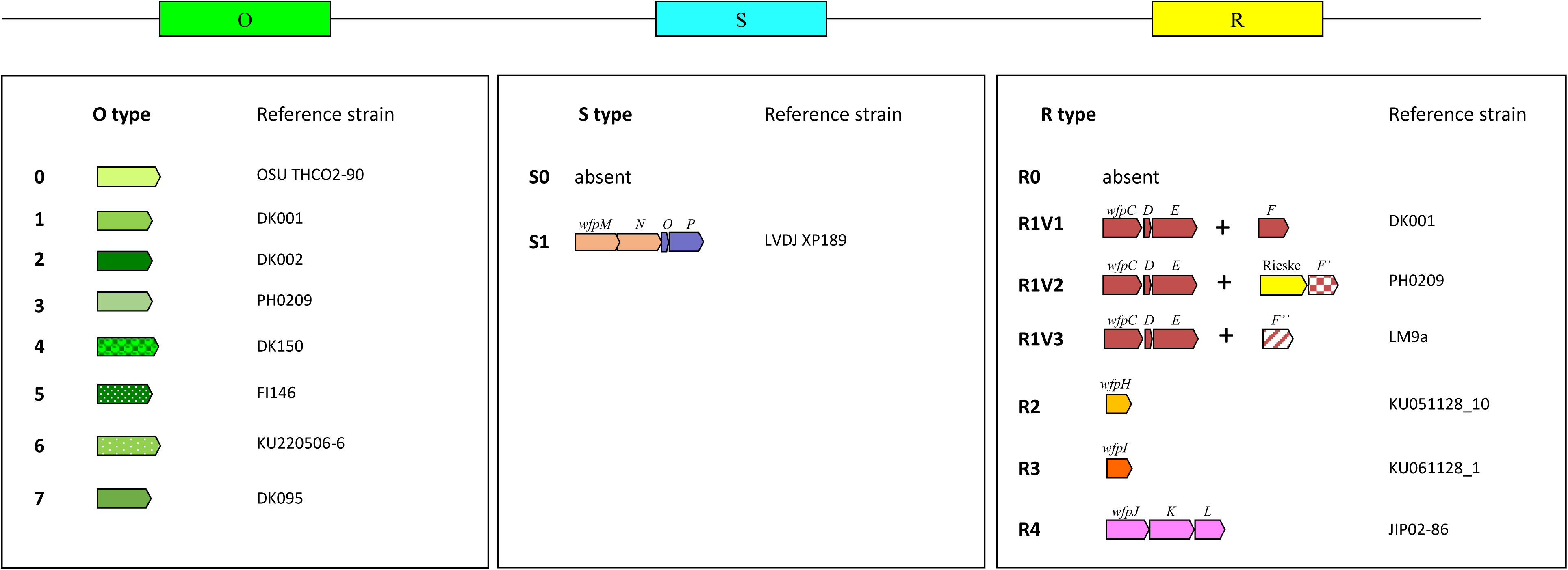
Three variable regions (O, S and R) of the O-Ag biosynthesis locus were identified in *F. psychrophilum*. The *in-silico* serotype is reported as the concatenation O:x–S:y–R:z.

The first set of biomarkers therefore rely on *wzy* (or *wzy*-like) genes and these biomarkers refer to O:x-types. Eight different O:x-types (O:0 to O:7) were recorded. Among genomes harboring *wzy1*, seven genomic architectures within the O-Ag biosynthesis locus were observed (hereafter O:1 subtypes) and for genomes harboring *wzy2*, three genomic arrangements were observed (hereafter O:2 subtypes). The selection of *wzy3* was obvious as it substitutes *wzy1/wzy2* in some strains and was accordingly chosen as a mPCR target gene (22). Among genomes harboring *wzy3*, five genomic architectures were observed (hereafter O:3 subtypes). The selection of the predicted *wzy4* followed the same principle as this gene lies next to *rmlA*, encodes as *wzy1, wzy2* and *wzy3* a multipass transmembrane protein and has been selected as a mPCR target (23). Among genomes harboring *wzy4*, three genomic architectures were observed (hereafter O:4 subtypes). The selection of additional *wzy-like* genes used a similar conceptual framework: located in the same gene cluster, encoding for a multipass transmembrane protein, unique and specific to a O-Ag genome group. Consequently, two genomic architectures were observed for *wzy5* (hereafter O:5 subtypes); three for *wzy6* (hereafter O:6 subtypes) and two for *wzy7* (hereafter O:7 subtypes). Finally, a unique genomic configuration (hereafter O:0 subtype) was observed for *wzy0* (nomenclature chosen according to Rochat and colleagues (22) and because this O:0 subtype encompasses the *F. psychrophilum* type strain). Because the *F. psychrophilum* Wzy proteins display poor sequence conservation, three-dimensional (3D) structure predictions and comparisons were used to identify remote similarities. Indeed, because protein 3D structures are typically more conserved than primary sequences, structure-based approaches provide a powerful means of identifying distantly related proteins and uncovering functional relationships that are not readily detectable from sequence comparison alone (43). Predicted template model (pTM) scores for individual Wzy proteins are reported in Supp. Table 4. All pTM scores were above 0.8, representing confident high-quality AlphaFold3 predictions. The DALI structural pairwise alignments scores (Z-score and rmsd) are presented in Supp Table 5.

Proteins encoded by THC0290_2043 (referring to Wzy0) and KU2205066_40100 (referring to Wzy6) displayed low (22.8% identity, 39.6% similarity) protein sequence conservation but similar 3D structures (Supp. Figure 2A) with significant metrics (Z score 31.8, rmsd 3.0). This was equally the case for Wzy0 and DK095_460107 (referring to Wzy7) displaying low (20.9% identity, 42.26% similarity) protein sequence conservation but superposable 3D structures (Supp. Figure 2B) with significant metrics (Z score 32.3, rmsd 2.7). To a lesser extent, DK150_v1_550174 (referring to Wzy4) displayed relevant 3D structural similarities to Wzy0 (Z score 17.5, rmsd 5.3), Wzy6 (Z score 14.4, rmsd 4.1) and Wz7 (Z score 19, rmsd 4.3). Significant structural conservation was also established for Wzy6 and Wzy7 (Z score 27, rmsd 3.4). Wzy2 and Wzy3, that also displayed moderate protein sequence conservation (33.7% identity, 50.1% similarity) possess 3D structures highly superposable (Supp. Figure 2C) with metrics of high significance (Z score 51.4, rmsd 1.7). Finally, Wzy1 and FI146_160075 (referring to Wzy5) did not display any significant structural similarities with any of the other Wzy proteins. Therefore, two groups of Wzy proteins had meaningful structural similarities: group 1 encompasses Wzy0, Wzy4, Wzy6 and Wzy7 whereas group 2 encompasses Wzy2 and Wzy3. Wzy1 and Wzy5 stand alone.

The second set of biomarkers relied on genes predicted to drive the biosynthesis and/or transfer of the R groups. This biomarker set refers to R:z types. R1, R2 and R3 groups have been previously reported by others (25). Genes predicted to be involved in the synthesis of an R4 group (and accordingly named *wfpJ, F* and *L*) substitute to *wfp* genes involved in R1, R2 and R3 group biosynthesis. Moreover, additional variations among the R1 group of genes were observed, allowing its split into three variants (*i.e.*, R1V1, R1V2 and R1V3).

A third set of biomarkers rely on 4 genes, located when present between *wbuA* and *flnA,* one of which encodes an acyl carrier protein and another for a beta-ketoacyl-[acyl-carrier-protein] synthase III family protein. The presence of these 4 genes was never observed linked to the R1 group but with R2, R3 and R4 instead. As the R1 group also requires these functions encoded by *wfpD* and *wfpC*, respectively, one might speculate that these 4 genes might be involved in the biosynthesis and/or transfer of an additional, different R’-group to a yet undetermined glycosidic residue of the O-Ag. We accordingly named these 4 genes *wfpM, N, O* and *P.* This biomarker set refers to S:y.

Supp. Figure 1 displays every single observed genetic structure and the bacterial strains selected as references for each. Table1 provides details regarding the selected genes chosen as biomarkers including bacterial strains selected as references and locus_tag of each gene or gene group. Therefore, the *in-silico* typing scheme relies on three sets of biomarkers leading to a combination composed of O:x-S:y-R:z where x = 0 to 7; S = 0 or 1 and R = 0 to 4 that further includes three R1 subdivisions (V1, V2 and V3) (Figure 2).

### Definition of the typing rules and thresholds

The thresholds applied by FlavoTyper to declare a marker present were derived empirically. Each biomarker sequence was searched with BLASTn against the reference genomes, and the percent identity and coverage of the best hit were recorded for every genome. The distribution of these values revealed two clearly separated populations: genomes carrying the marker produced near-identical and near-complete alignments, whereas genomes lacking it produced either divergent or only partially aligned hits. After examination of each marker, the threshold was set at the lowest identity and coverage values reached by genomes carrying the marker; placed at this boundary, the retained thresholds (97% identity and 94% coverage) discriminate between genuine hits from partial or false ones. The per-marker distributions for each biomarker are provided in Supp. Figure 3.

**Figure 3:**
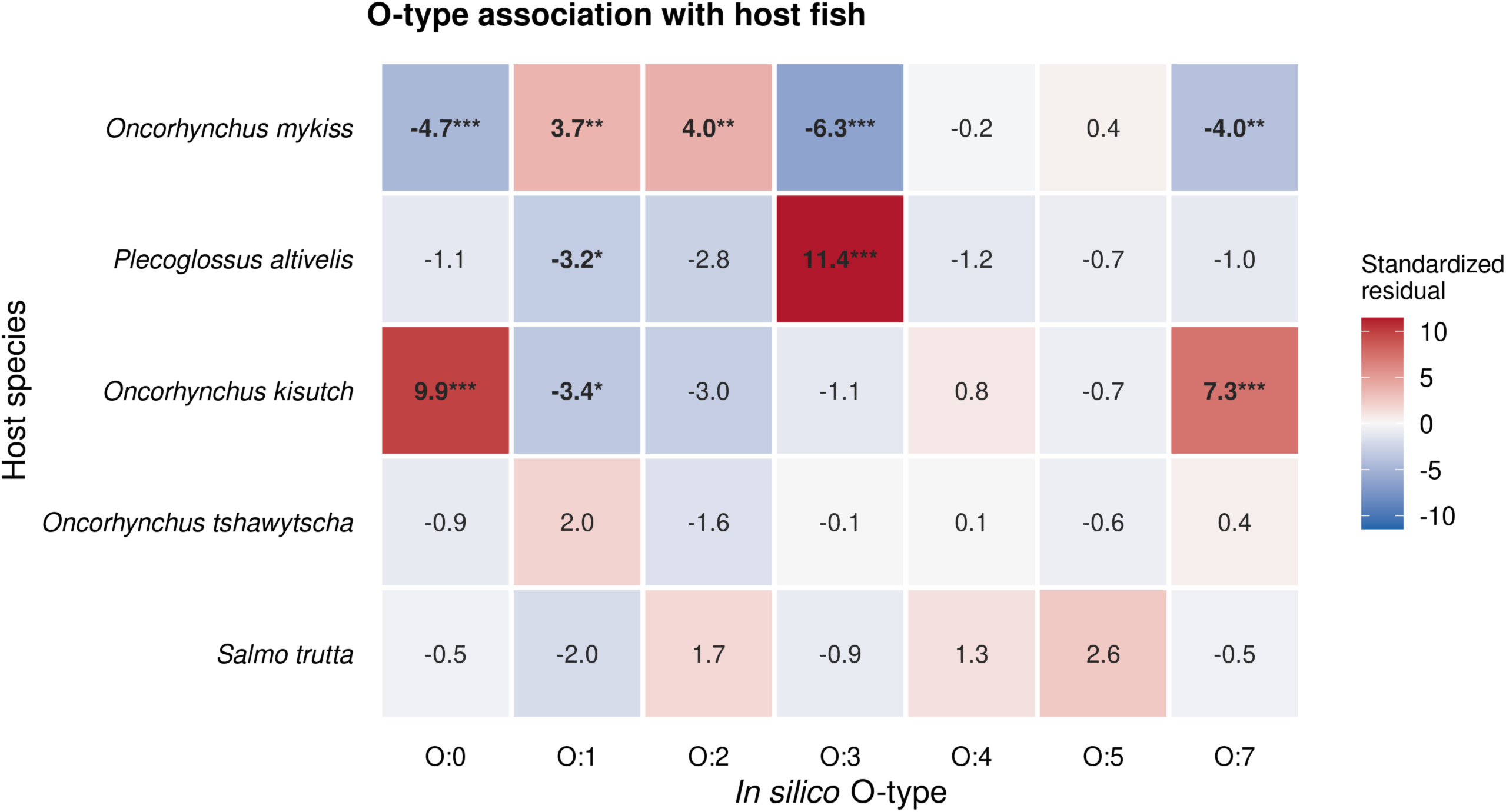
Standardized Pearson residuals for in silico O-type by host fish. Red = enriched, blue = depleted; cell values are the residuals, with Holm-adjusted significance (* p < 0.05, ** p < 0.01, *** p < 0.001).

### Application to the NCBI repository genome dataset

Each of the 482 *F. psychrophilum* assemblies, publicly available at the time of writing, and retrieved from the NCBI repository was processed with FlavoTyper using the default parameters (species-verification threshold ANI ≥ 95 % against the type strain NCIMB 1947ᵀ; marker thresholds of ≥ 97 % identity and ≥ 94 % coverage). Out of the 482 genomes, 407 (84.4 %) passed the species validation step and were typed, while 75 (15.6 %) were rejected at the quality-control step and reported as NotTyped. Among the 407 typed genomes, 375 (77.8 % of the dataset) received a fully resolved serotype; 9 (1.9 %) a partial call; 4 (0.8 %) an ambiguous call; and 19 (3.9 %) an undefined call (no serotype-defining marker detected). Across the 375 resolved genomes, FlavoTyper identified 23 distinct serotypes in total, whose frequencies are provided in Supp. table 6. The two serotypes dominating the dataset are O:2–S0–R1V1 (n = 134) and O:1–S0–R1V1 (n = 69), together accounting for more than half of the resolved genomes, followed by O:1–S0–R4 (n = 43) and O:0–S0–R0 (n = 21); the remaining serotypes were each represented by twenty or fewer genomes.

The 75 genomes excluded by the quality-control step were examined (Supp. table 7). The computed ANI for those assemblies was below the 95 % species threshold: 76.4–78.5 %, median 77.4 %; for 31 of them fastANI returned no hits at all. It is in sharp contrast with the 407 typed genomes, whose ANI ranged from 98.7 % to 100.0 % (median 99.1 %). The rejected assemblies (*i.e.*, those that did not pass the 95 % species delineation threshold) do not belong to *F. psychrophilum* species in full concordance with the NCBI taxonomy check (*i.e.*, “Failed” or an “Inconclusive” status). Partial and undefined calls typically arise from highly fragmented genomes and/or assembly issues; whereas ambiguous calls arise from genomes sequenced from multiple strains (not pure culture). Together, these results demonstrate that FlavoTyper scales to the full public collection of *F. psychrophilum* genome assemblies, recovering 23 distinct serotypes. Importantly, the species-verification step reliably prevents assemblies from misclassified species from being assigned a spurious serotype whereas partial, ambiguous and undefined calls point out different issues that may require additional inspections and/or cross-checks.

### Links with host fish species

The *F. psychrophilum* PubMLST genome collection has been manually curated and importantly does not contain duplicated stains. *In-silico* serotyping using this genome collection allowed the assignment of a resolved serotype (*i.e.*, O:x-S:y-R:y) to 334 out of the 357 assemblies publicly available at the time of writing (i.e., 94%). For the remaining 23 genomes: 7 were partial; 2 ambiguous; 14 undefined; and none were NotTyped. The full PubMLST isolates collection, together with the associated metadata and the serotypes assigned by FlavoTyper, is provided in Supp. table 8.

Because this dataset is rather small, we only focus on the O:x-types not taking into account additional variants (*i.e.*, S:y-R:y types). These O:x-types were strongly structured by host fish species (Fisher’s exact test, p < 10⁻⁵ on the restricted set of n = 302). Post-hoc examination of the standardized residuals revealed several strong, biologically coherent associations (Figure 3). Ayu (*Plecoglossus altivelis*) was almost exclusively associated with O:3 (residual +11.4; 20/24 isolates); coho salmon (*O. kisutch*) was enriched for O:0 (+9.9) and O:7 (+7.3) and depleted of O:1; and rainbow trout (*O. mykiss*) was enriched for O:1 (+3.7) and O:2 (+4.0) and depleted of O:0, O:3 and O:7. No significant association was detected for *O. tshawytscha* or *Salmo trutta*. These associations reproduce a host–serotype relationship documented by the previously developed serological and mPCR-based typing methods (5,17,20,22). These association trends are likely functional rather than merely distributed. One can hypothesize that these links provide the bacterium a fitness advantage because serotypes O:1 and O:2 (corresponding to Fd and Th serotypes, respectively) are predominant in rainbow trout isolates causing disease outbreaks (20,44,45). In addition, these O:1 and O:2 isolates tend to be more virulent than others in rainbow trout experimental infection models (46,47,48). On the other hand, O:3 isolates are predominant and virulent to ayu (49), whereas O:0 isolates are predominant and virulent to coho salmon (50) suggesting that the O-Ag on his own contributes to host specificity and/or virulence rather than acting as a passive marker.

### Impact of recombination: O-Ag locus shuffling

The distribution of *in-silico* resolved (O:x-S:y-R:y) serotypes showed limited concordance with the core genome-based phylogeny, with marked incongruence between biomarker groups and phylogenetic relationships (Figure 4). Rather than segregating with particular lineages, the O-types were dispersed across the tentative tree, and most clonal complexes were polymorphic for O-type. The largest lineage, CC-ST10, encompassed alone five distinct O-types (O:1, O:2, O:3, O:5 and O:6), with O:2 and O:1 predominating; CC-ST90 likewise combined O:1, O:2 and O:3, and even the CC-ST56 (seven isolates only) spanned four different O-types. Conversely, individual O-types recurred in phylogenetically unrelated backgrounds: O:1 was found in five clonal complexes (CC-ST10, CC-ST90, CC-ST52, CC-ST56 and CC-ST124) and O:3 also in five (CC-ST10, CC-ST52, CC-ST56, CC-ST90 and CC-ST191). This pattern of closely related strains displaying different serotypes cannot be explained by vertical inheritance and instead points to frequent horizontal gene exchange arising from homologous recombination within the O-Ag biosynthesis locus. High recombination is a hallmark of the evolutionary process of the species with a (r/m) estimated to ∼13 (51). It has been previously reported that highly similar strains (*e.g.*, FI056 and DK002) differed almost exclusively at the polysaccharide-biosynthesis locus, through mutually substituting genes occupying the same chromosomal position (22). The S- and R-biomarkers similarly showed a dispersed distribution across the phylogenetic tree. Together, these results indicate that the O-Ag locus behaves as a recombination hotspot, shuffled among lineages independently of the genomic backbone. This decoupling was not absolute as O:0 was essentially restricted to CC-ST09, showing that some serotype–lineage associations remain “clonally” conserved and that O-Ag exchange, though pervasive, is not uniform across the population.

**Figure 4:**
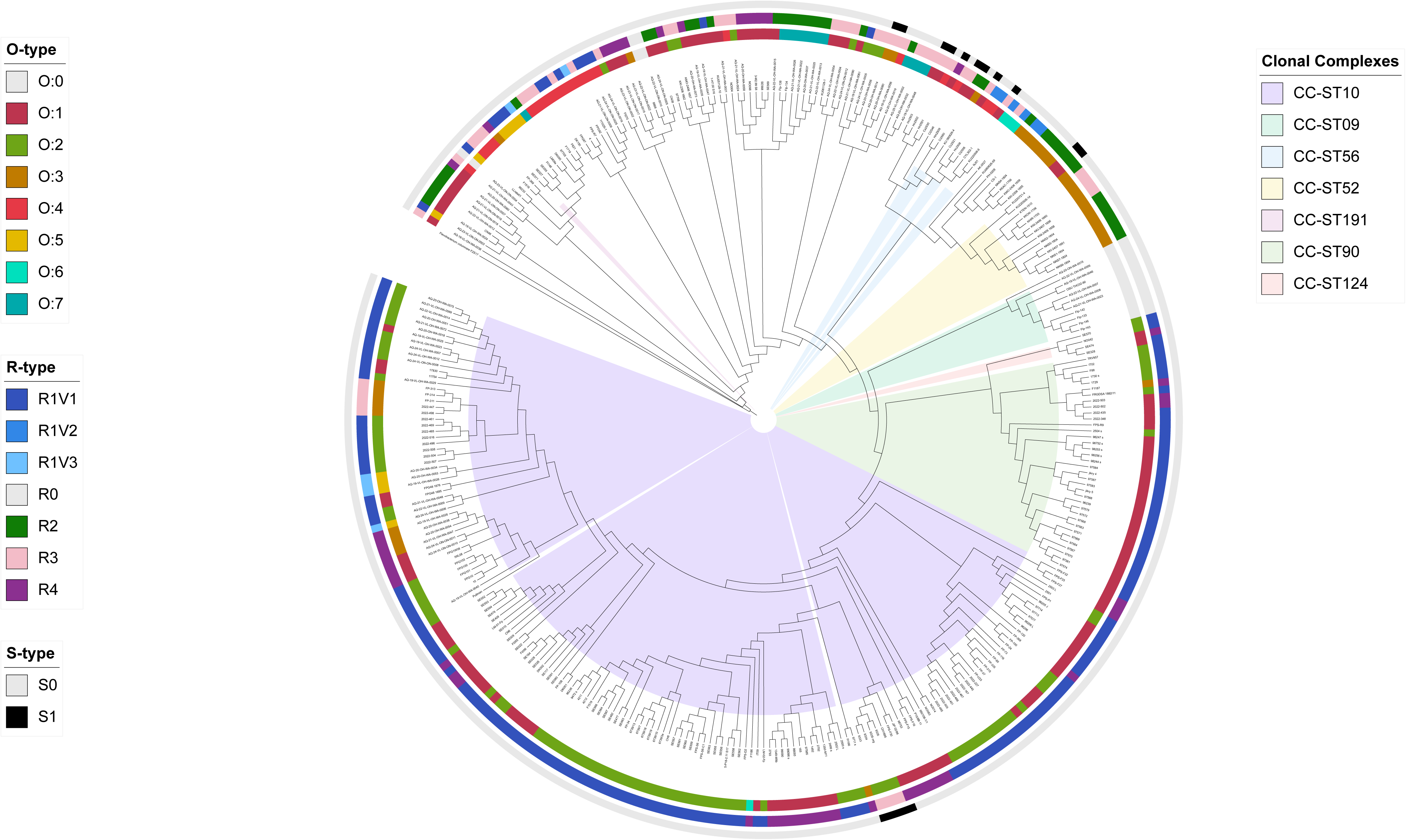
*In-silico* serotypes and evolutionary relationships between 334 *F. psychrophilum* genomes. The O-, R- and S-types are shown on a tree capturing the phylogenetic relationships between the 334 *F. psychrophilum* strains retrieved from PubMLST. The tree was constructed by maximum-likelihood inference from a concatenated alignment of 869 single-copy core protein families (254,073 amino-acid positions; see Materials and Methods for details) and rooted on *F. columnare* strain F2S17 as an outgroup. The three concentric rings indicate, from inner to outer, the O-, R- and S-types of each strain. Clonal complexes were retrieved from PubMLST.

## Conclusion

As WGS becomes increasingly established as a routine approach, robust *in-silico* serotype prediction methods are essential for ensuring reliable and standardized typing. FlavoTyper was specifically developed to meet this goal. The output is designed for straightforward interpretation and includes explicit quality-control metrics to facilitate serotyping. In addition, the FlavoTyper marker database could be easily upgraded if yet unobserved biomarkers are identified in the future.

We believe that this tool will be useful for a better understanding of host-pathogen interaction, including host specificity. Indeed, the recent expansion of *F. psychrophilum* in non-salmonid fish including eel (7) and carp (8,9,52) will require robust approaches for epidemiological surveillance and for tracking disease outbreaks. This tool has also been implemented in PubMLST (29) in order to enable simultaneous sequence typing and serotype predictions from WGS data. Beyond academic research, this framework could have important practical applications, including relevant bacterial strain selection for vaccine development and selective breeding for disease resistance.

## Author statements

### Author contributions

Author contributions following the CRediT taxonomy are as follows. SM developed software, performed formal analyses and visualizations; PD, MC and TR contributed to the methodology and conducted the investigations; KJ and NH developed software and performed formal analyses; ED contributed to the conceptualization and methodology, acquired funding, administered the project and wrote the original draft of the manuscript. All authors read and approved the final manuscript.

### Consent for publication

No ethical or consent-related approval was required for the research in this study.

## Acknowledgements

We are grateful to Valentin Loux (INRAE, MaIAGE, France) for software testing, feedback and helpful discussions and to Dr. Erina Fujiwara-Nagata (Kindai University, Japan) for providing genomic sequences of additional strains. The LABGeM (CEA/Genoscope & CNRS UMR8030), the France Génomique and French Bioinformatics Institute national infrastructures (funded as part of Investissement d’Avenir program managed by Agence Nationale pour la Recherche, contracts ANR-10-INBS-09, ANR-11-INBS-0013 and ANR-21-ESRE-0048) are acknowledged for support within the MicroScope annotation platform.

