## Supplemental figures for "FlavoTyper: a genome-based *in-silico* serotyping tool for the fish pathogen *Flavobacterium psychrophilum*"

**Supplementary Figure 1:** Genomic organization of the *F. psychrophilum* O-Ag encoding loci. The O-Ag encoding loci were analyzed using the Microscope platform. Gene conservation was predicted using both homology and synteny criteria. When available gene names have been included, GT denotes putative glycosyl transferases. They have been colored-coded according to their predicted functional involvement in O-Ag biosynthesis and/or putative function (*e.g*, L-Rha, L-FucNAc and the D-Qui*2NAc* synthesis and transfer, respectively.) S1 and R groups have been underlined. Sub-types were grouped according to the *wzy* (or *wzy*-like) gene, denoted by an arrow. All observed sub-types are reported with their corresponding biomarkers combination $O_x$-$S_y$-$R_z$ from selected strains chosen as references. At the time of writing and analyzing all available genomes retrieved from Genbank, a unique sub-type was observed in O:0; seven for O-1; three for O:2; five for O:3; three for O:4; two for O:5; three for O:6 and two for O:7.

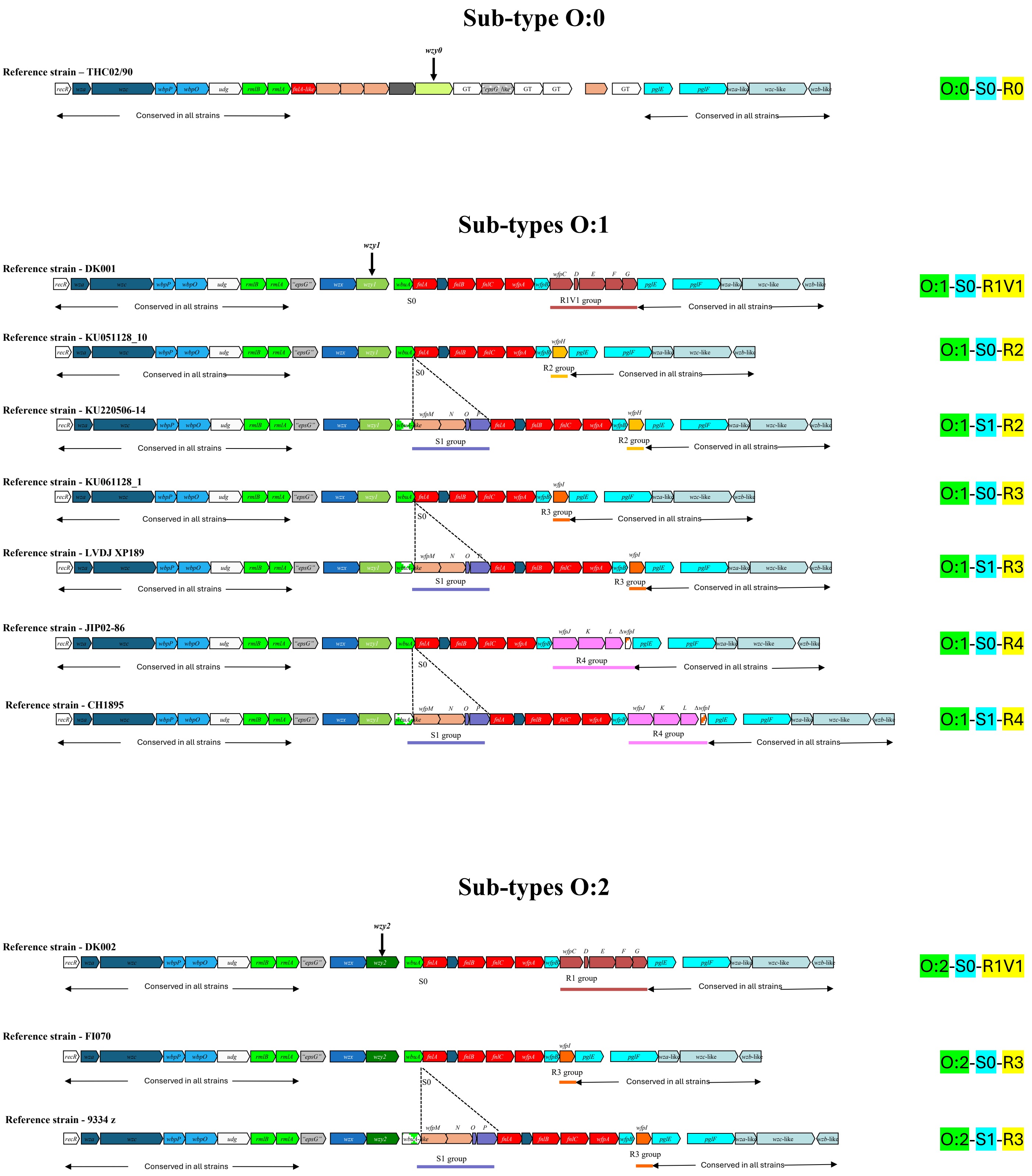

### Sub-types O:3

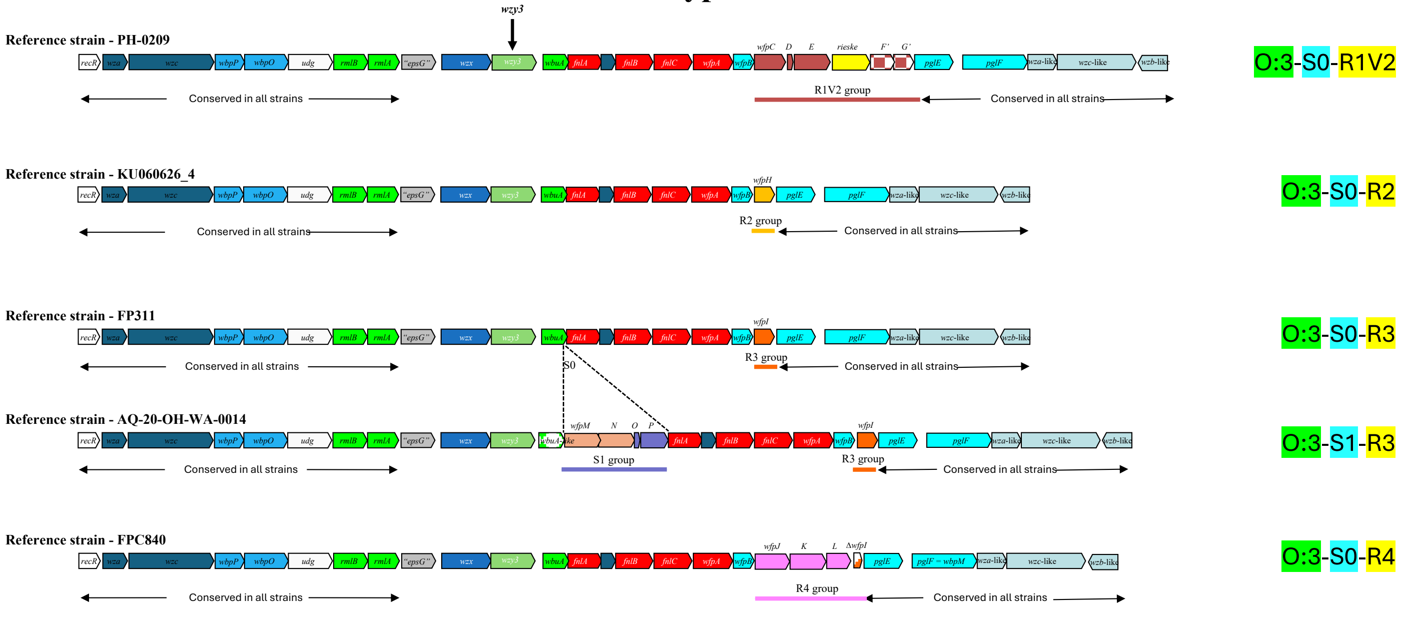


### Sub-types O:4

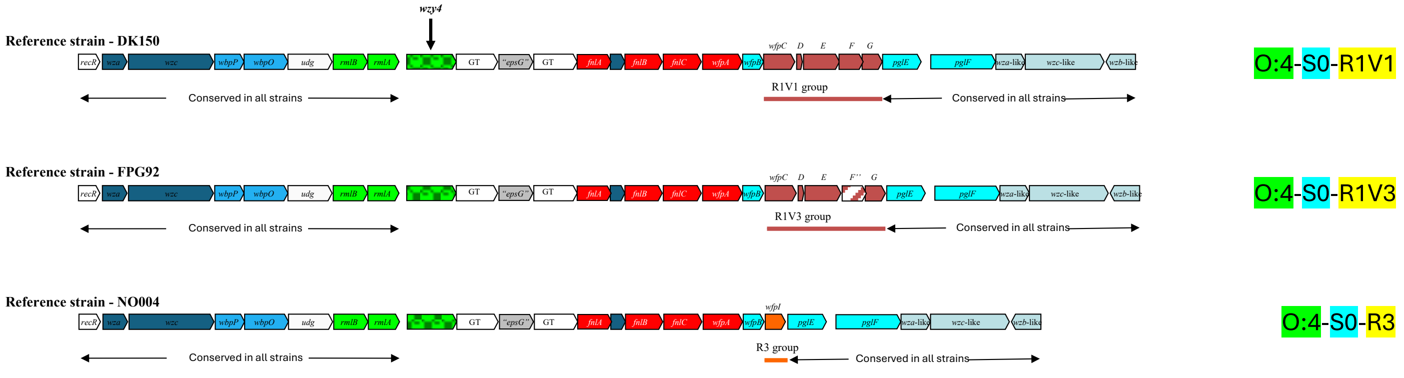


### Sub-types O:5

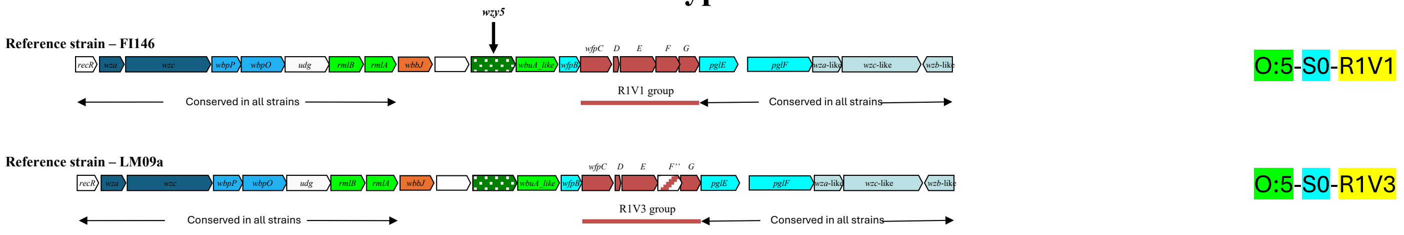


### Sub-types O:6

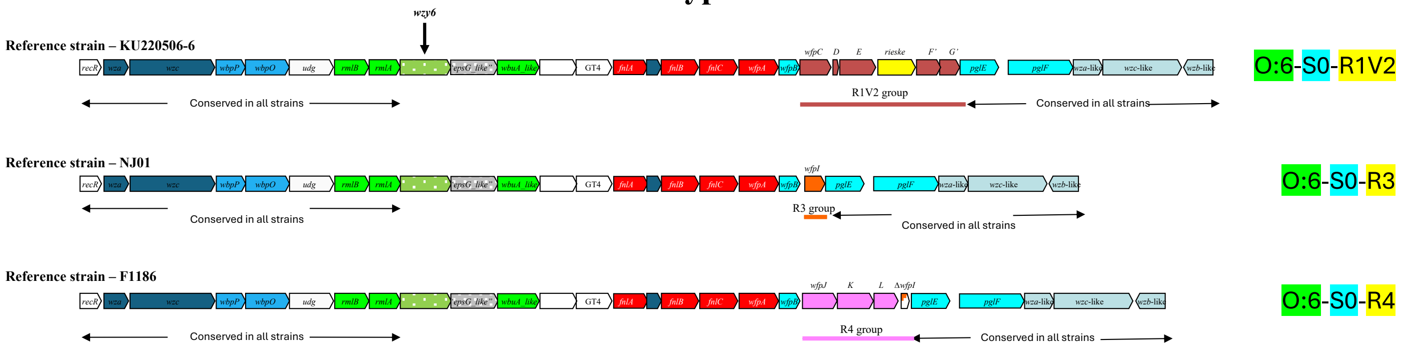


### Sub-types O:7

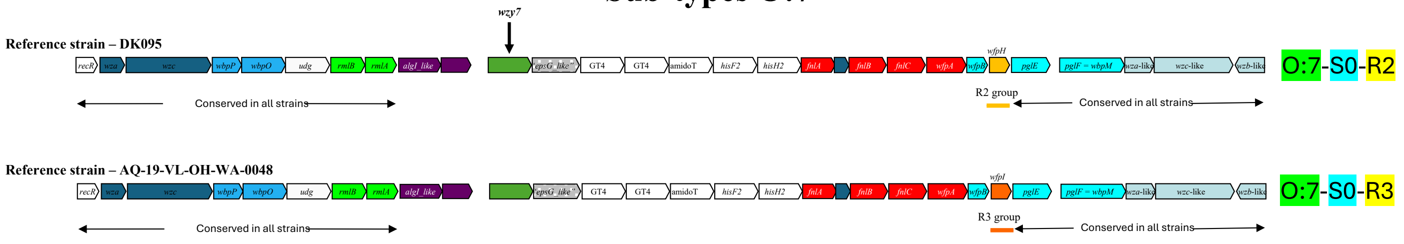

**Supplementary Figure 2:** Structural comparisons of selected *F. psychrophilum* Wzy proteins retained as O:x type biomarkers predicted with AlphaFold (V3). Left panel, top view and right panel side view. A) THC0290_2043 (Wzy0) in blue and KU2205066_40100 (Wzy6) in yellow B) THC0290_2043 (Wzy0) in blue and DK095_460107 (Wzy7) in yellow and C) DK002_320117 (Wzy2) in blue superposed with PH0209_2216 (Wzy3) in yellow.

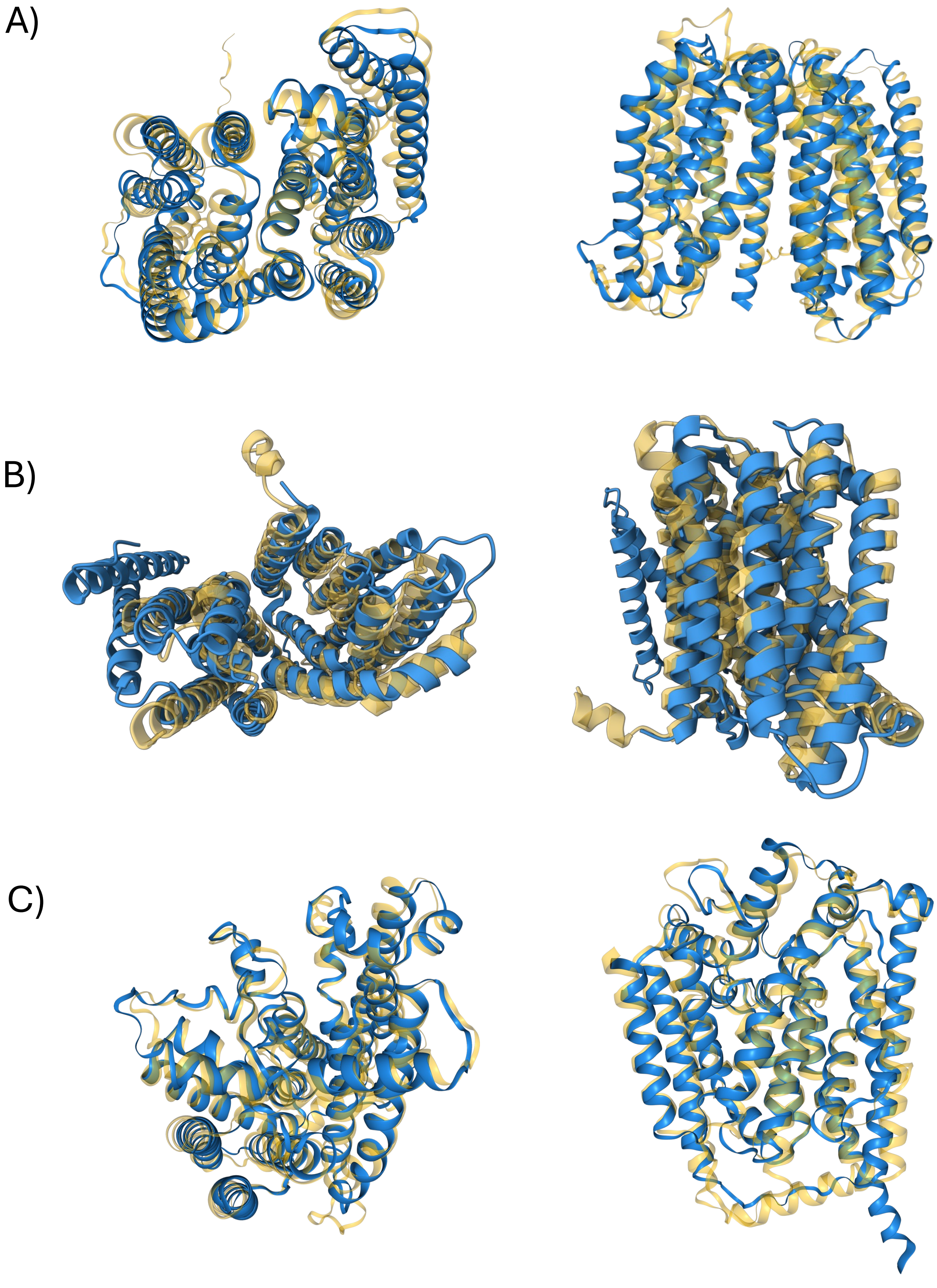

**Supplementary Figure 3:** Per-marker distribution metrics.
Distribution of the BLASTn percent identity (left panel) and percent coverage (right panel) values for selected biomarkers across the reference genomes. Each point represents one genome; blue points denote genomes that possess the marker and orange points, genomes that lack it. Proteins selected as A) O-type biomarkers; B) S-type biomarkers and C) R-type biomarkers.

A)

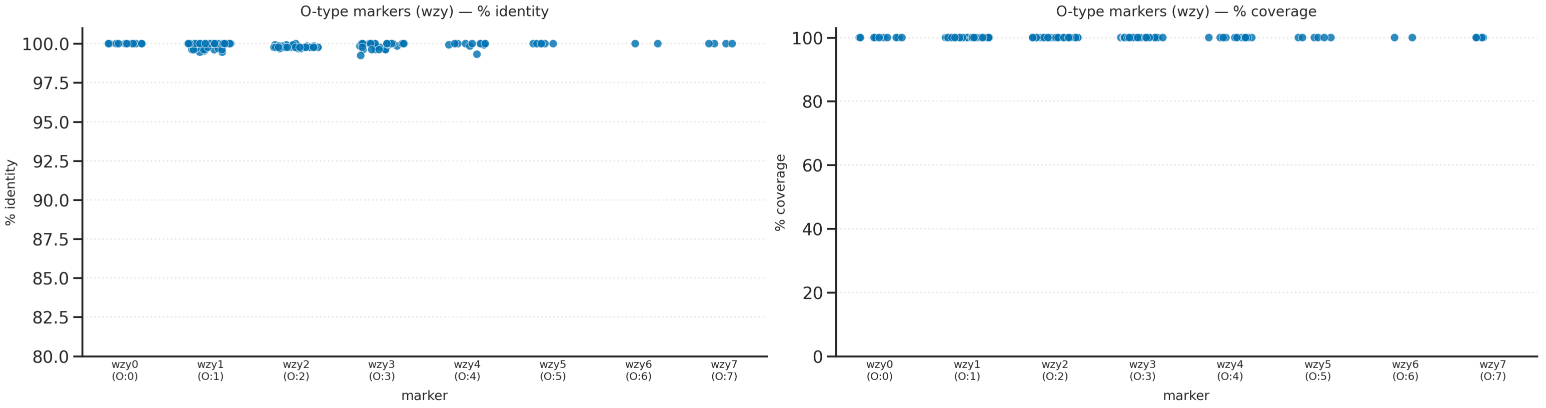


B)

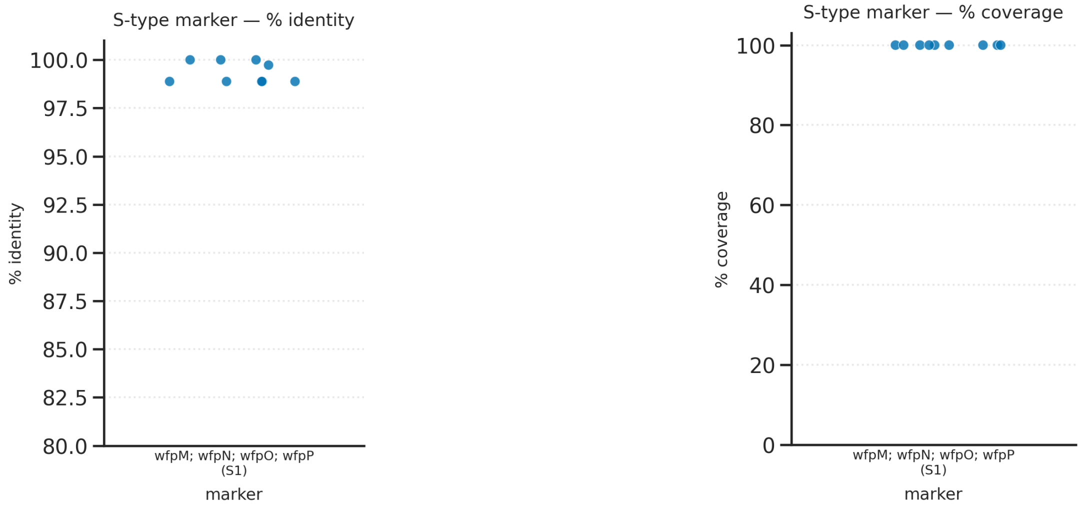


C)

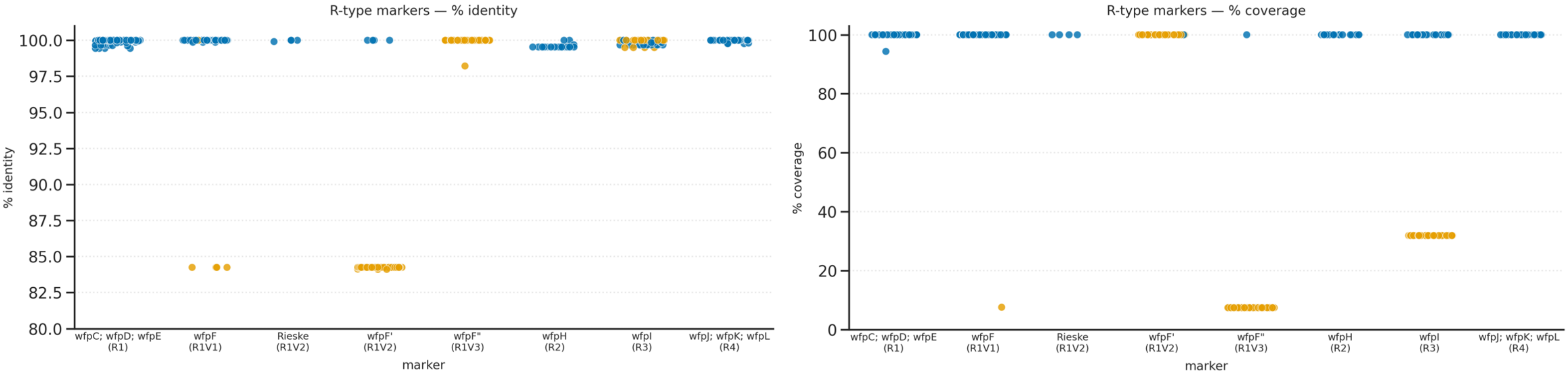
